# Analysis of human GPER expression in normal tissues and select cancers using immunohistochemistry

**DOI:** 10.1101/2023.03.09.531931

**Authors:** Christopher A. Natale, John T. Seykora, Todd W Ridky

**Affiliations:** Linnaeus Therapeutics, Haddonfield, New Jersey, 08033; Perelman School of Medicine, Department of Dermatology, University of Pennsylvania, Philadelphia, Pennsylvania 19104

## Abstract

GPER (G protein-coupled estrogen receptor) has been reported to play roles in several areas of physiology including cancer, metabolic disorders, and cardiovascular disease. However, the understanding of where this receptor is expressed in human tissue is limited due to limited available tools and methodologies that can reliably detect GPER protein. Recently, a highly specific monoclonal antibody against GPER (20H15L21) was developed and is suitable for immunohistochemistry. Using this antibody, we show that GPER protein expression varies markedly between normal human tissue, and also among cancer tissue. As GPER is an emerging therapeutic target for cancer and other diseases, this new understanding of GPER distribution will likely be helpful in design and interpretation of ongoing and future GPER research.

## Introduction

GPER (G protein-coupled estrogen receptor) is a transmembrane protein that is activated by endogenous estrogens and has been reported to play a role in regulating various physiological processes. Unlike the classical nuclear estrogen receptors, GPER does not require a genomic mechanism for its actions and can directly activate G protein-coupled signaling pathways (1). This direct activation results in rapid and localized responses to estrogens. GPER is thought to be primarily expressed in the endothelial cells of blood vessels and in the hypothalamus and pituitary gland of the brain (2). GPER has been implicated in various diseases, including cancer, metabolic disorders, and cardiovascular disease, making it a target for drug development (3,4).

G protein-coupled receptors (GPCRs) are challenging targets for antibody development due to their dynamic structures, low expression levels, conserved epitopes, and structural variability. These properties make the development of specific antibodies difficult (5). Additionally, since GPCRs typically have low levels of RNA expression and RNA expression does not correlate with protein levels (6), the understanding of where GPER is expressed in human tissues remains limited.

Recently, a novel rabbit monoclonal anti-human GPER antibody, 20H15L21, was identified (7). The specificity of this antibody clone for GPER was determined using western blot analyses, immunofluorescence, and genetic manipulation of GPER. A series of normal and cancer formalin-fixed, paraffin-embedded (FFPE) human tissue samples were assessed using immunohistochemistry. GPER was found to be expressed in several normal tissues, including brain, pancreas, stomach, small intestine, liver, kidney, adrenal glands, and placenta. Additionally, GPER was found to be expressed in several types of cancer including hepatocellular, pancreatic, renal, and endometrial cancers, pancreatic neuroendocrine tumors, and pheochromocytomas.

Here, we have an expanded analysis on FFPE human tissues to further assess where GPER is expressed in normal tissues and select malignancies. We have performed immunohistochemistry using the 20H15L21 GPER antibody on 33 distinct normal human tissue samples as well as >30 cases of colon, pancreatic, cutaneous melanoma, lung, prostate, and breast human cancer tissue.

## Methods

### Tissue Microarrays

FFPE tissue microarrays were purchased from Biomax (Derwood, MD, USA). Catalog numbers for the tissue arrays are as follows: Normal tissue (FDA999y2), Melanoma (ME1002b), and multiple other cancers (BC000119b).

### GPER Immunohistochemistry and Imaging

Tissue microarrays were stained using the GPER recombinant rabbit monoclonal antibody clone:20H15L21 (Thermo Fisher Scientific, catalog #703480) by DCL Pathology (Indianapolis, IN, USA). IHC staining for GPER was performed using the Ventana Benchmark Ultra platform (Roche Diagnostics). Tissue sections were cut onto positively charged slides at a thickness of 3-5 microns and baked at 60-65 degrees Celsius for 10 minutes prior to IHC staining. Slides were deparaffinized on the Benchmark Ultra at 72 degrees Celsius using EZ Prep solution (Cat. No. 950-102). Slides were then treated with Ultra Cell Conditioning Solution 1 (950-224) for 36 minutes at 95 degrees Celsius. The antibody used for this assay was anti-GPER (20H15L21) rabbit monoclonal antibody and was diluted 1:375 in Ventana Antibody Diluent (Cat. No. 251-018). 100 μL of diluted antibody was applied to the slides and incubated for 32 minutes at 37 degrees Celsius. After the primary antibody incubation, the slides were rinsed and detection was performed using the Roche ultraView Universal Alkaline Phosphatase Red Detection Kit (Cat. No. 760-501). After the detection reaction was completed, tissues were counterstained with Hematoxylin II (Cat. No. 790-2008) for 4 minutes and blued with Bluing Reagent (Cat. No. 760-2037) for 4 minutes. Slides were then rinsed with soapy water followed by DI water and dehydrated in 100% reagent alcohol. Slides were transferred to xylene and coverslipped. Stained slides were Whole Slide Imaged using the Motic EasyScanPro platform and stored as Leica Aperio SVS files.

### Quantification

Scanned slide images were viewed with QuPath (8) and images of the entire core were exported for all evaluable, non-defective cores. Core images were then assessed for the extent of GPER staining by calculating the area of GPER staining using Adobe Photoshop. The total area of the core was measured in pixels, subtracting any acellular regions representing >10% of the core area. The area of GPER staining was measured in pixels using a color range that captures red staining indicative of GPER protein expression. The area of GPER staining was divided by the cellularized area of the tissue core and multiplied by 100, resulting in a percent GPER staining metric. Tissue staining results were reviewed by a board-certified pathologist adjusted when necessary to account for staining in the normal or cancer tissue of interest. All quantification data is in Supplementary Data Table 1.

## Results

The extent of GPER protein expression was assessed in 33 distinct normal tissues, with 3 cases for each tissue type (Figure 1). The normal tissue types with >50% GPER protein expression on average were cerebrum, tonsil (epithelium), liver, small intestine (epithelium), kidney, pituitary gland, colon (epithelium).

**Figure 1:**
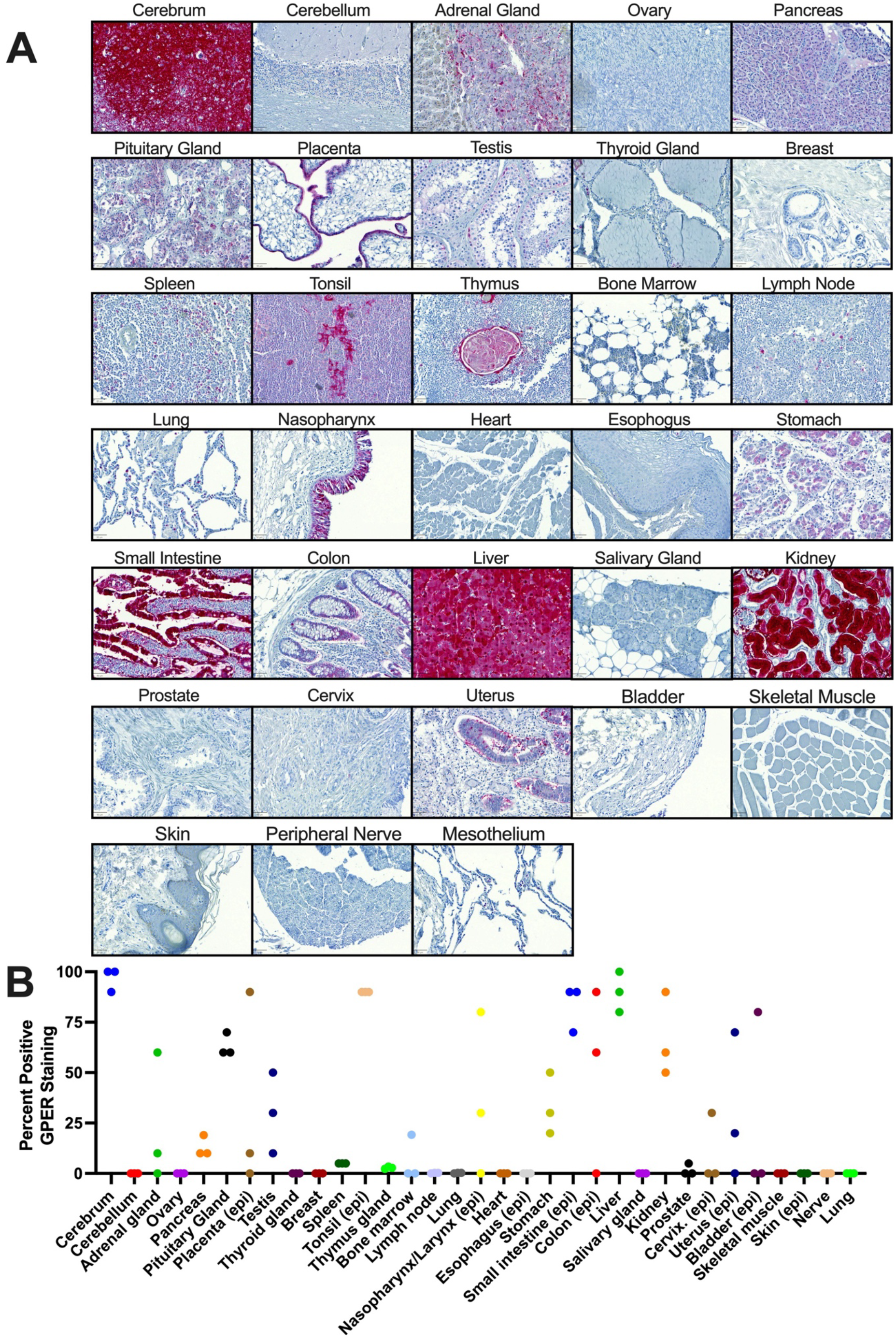
GPER staining in normal tissues. (A) Representative images of GPER+ areas in normal tissues, scale bar = 50μm. (B) Quantification of GPER staining across normal tissues, epithelial tissues were quantitated with staining in the epithelial region of the tissue specifically, and denoted with (epi).

The extent of GPER protein expression was also assessed in 6 different cancer types, with more than 30 cases for each cancer type (Figure 2). GPER was most highly expressed in colon cancer cases, followed by pancreatic cancer, melanoma and lung cancer. GPER was expressed in a low percentage of prostate and breast cancer cases, with 2/36 and 0/31 cases, respectively, having >1% GPER stained area.

**Figure 2:**
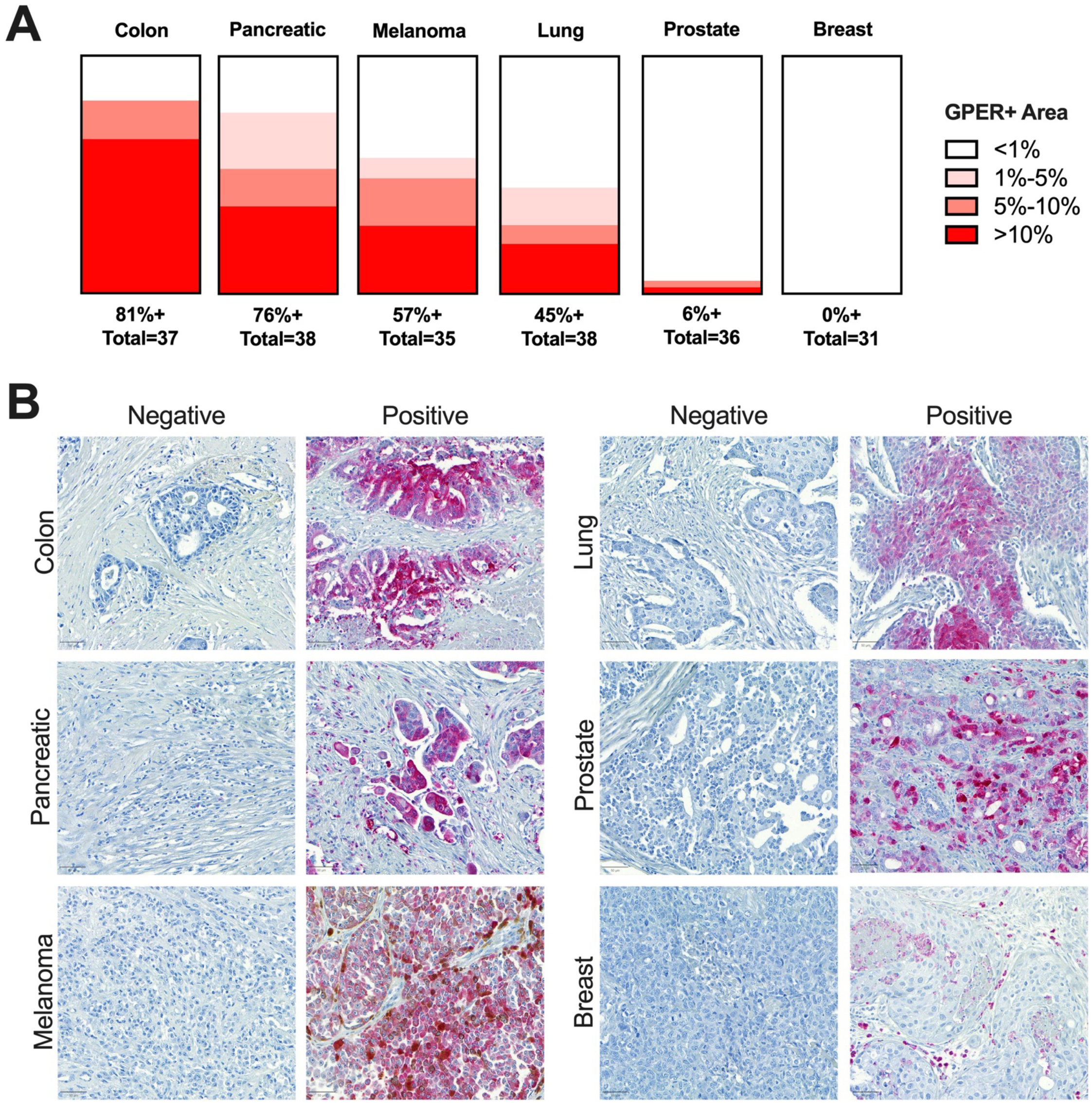
GPER staining in select malignancies. (A) Visualization of GPER staining across colon, pancreatic, melanoma, lung, prostate, and breast cancer cases. (B) Representative images of negative and positive GPER staining in the examined cancer types, scale bar = 50μm.

Although adjacent normal and cancer tissue could not be assessed in these samples, there appears to be concordance of GPER protein expression between normal and cancer tissues. For example, normal colon and pancreas tissue expresses GPER protein, and a high percentage of colon and pancreatic cancer cases also expresses GPER protein. Conversely, we observed low GPER protein expression in both normal breast and prostate, and a similarly low GPER protein expression in breast and prostate cancer cases.

## Discussion

Here we described GPER protein expression in 33 normal tissue types and 6 common cancer types. The antibody labeling was robust in the IHC application demonstrating very low background and a wide dynamic range, labeling GPER in both cytoplasm and the plasma membrane. Our findings are consistent with and validate the initial report of the monoclonal anti-human GPER antibody, clone 20H15L21(7). In the prior and current study, GPER expression was observed in many of the same normal tissues; including cerebrum, pancreas, liver, small intestine, placenta, and kidney. While the prior study assessed a larger number of cancer types with a smaller number of cases for each, this study found that GPER protein was expressed in the high percentage of colon, pancreatic, melanoma, and lung cancer cases, consistent with reports from preclinical models (9–13). While GPER expression was detected in a high percentage of these cancer types, there is heterogeneity that exists between cases of the same cancer type, suggesting that GPER protein expression in patient samples may be a useful biomarker for GPER-directed therapies that are in development, like LNS8801 (14,15).

Additionally, both studies found a low percentage of prostate cancer samples express GPER and rarely, if at all, in breast cancer samples. The observation that GPER protein is rarely expressed in human breast cancer cases is noteworthy, given the substantial number of reports discussing the role of GPER in preclinical models of breast cancer (16). Although GPER signaling was originally reported to be tumor promoting in some breast cancer models (17), subsequent reports show that GPER signaling inhibits breast cancer (18–20). Additionally, several studies have documented correlations between GPER expression and prognosis in breast cancer, with contradictory results (21,22). While there certainly may be correlations between GPER expression and clinical features of certain cancer types, this approach does not address whether or not GPER is signaling actively, thus interpretation of the results is limited. Given the lack GPER protein expression in normal or cancer human breast tissue, it is unclear whether GPER plays a significant role in this tissue type.

Given the wide range of GPER-mediated functions that have been identified in various tissue types, assays and reagents that can reliably detect GPER protein in human tissues are critically important to further the understanding of GPER biology. Based on the compelling initial report of the monoclonal anti-human GPER antibody, clone 20H15L21 (7), we have conducted an expanded analysis on FFPE human tissues to further document GPER distribution in normal tissues and select malignancies. The monoclonal anti-human GPER antibody, clone 20H15L21 is an important tool that will enable the scientific community to determine GPER protein in human FFPE tissues.

## Acknowledgments

The authors would like to thank Justine Cohen, Jordi Rodon Ahnert, Patrick Mooney, and Tina Garyantes for their critical pre-submission review of this manuscript. The authors would also like to thank Bryan Szpunar for overseeing this project at DCL Pathology.

## References

1. Prossnitz ER, Arterburn JB. International Union of Basic and Clinical Pharmacology. XCVII. G Protein–Coupled Estrogen Receptor and Its Pharmacologic Modulators. Pharmacol Rev [Internet]. 2015 Jul 1 [cited 2023 Jan 29];67(3):505. Available from: /pmc/articles/PMC4485017/

2. Isensee J, Meoli L, Zazzu V, Nabzdyk C, Witt H, Soewarto D, et al. Expression pattern of G protein-coupled receptor 30 in LacZ reporter mice. Endocrinology. 2009 Apr;150(4):1722–30.

3. Arterburn JB, Prossnitz ER. G Protein–Coupled Estrogen Receptor GPER: Molecular Pharmacology and Therapeutic Applications. https://doi.org/101146/annurev-pharmtox-031122-121944 [Internet]. 2023 Jan 20 [cited 2023 Jan 30];63(1):295–320. Available from: https://www.annualreviews.org/doi/abs/10.1146/annurev-pharmtox-031122-121944

4. Prossnitz ER, Barton M. The G-protein-coupled estrogen receptor GPER in health and disease. Nat Rev Endocrinol [Internet]. 2011;7(12):715–26. Available from: https://www.ncbi.nlm.nih.gov/pubmed/21844907

5. Hutchings CJ, Koglin M, Olson WC, Marshall FH. Opportunities for therapeutic antibodies directed at G-protein-coupled receptors. Nature Reviews Drug Discovery 2017 16:11 [Internet]. 2017 Jul 14 [cited 2023 Jan 30];16(11):787–810. Available from: https://www.nature.com/articles/nrd.2017.91

6. Schiöth HB, Nordström KJV, Fredriksson R. Mining the gene repertoire and ESTs for G protein-coupled receptors with evolutionary perspective. Acta Physiologica [Internet]. 2007 May 1 [cited 2023 Jan 30];190(1):21–31. Available from: https://onlinelibrary.wiley.com/doi/full/10.1111/j.1365-201X.2007.01694.x

7. Bubb M, Beyer ASL, Dasgupta P, Kaemmerer D, Sänger J, Evert K, et al. Assessment of G Protein-Coupled Oestrogen Receptor Expression in Normal and Neoplastic Human Tissues Using a Novel Rabbit Monoclonal Antibody. Int J Mol Sci [Internet]. 2022 May 1 [cited 2023 Jan 30];23(9). Available from: /pmc/articles/PMC9099907/

8. Bankhead P, Loughrey MB, Fernández JA, Dombrowski Y, McArt DG, Dunne PD, et al. QuPath: Open source software for digital pathology image analysis. Scientific Reports 2017 7:1 [Internet]. 2017 Dec 4 [cited 2023 Jan 30];7(1):1–7. Available from: https://www.nature.com/articles/s41598-017-17204-5

9. Natale CA, Li J, Zhang J, Dahal A, Dentchev T, Stanger BZ, et al. Activation of G protein-coupled estrogen receptor signaling inhibits melanoma and improves response to immune checkpoint blockade. Elife [Internet]. 2018;7. Available from: https://www.ncbi.nlm.nih.gov/pubmed/29336307

10. Natale CA, Li J, Pitarresi JR, Norgard RJ, Dentchev T, Capell BC, et al. Pharmacologic Activation of the G Protein–Coupled Estrogen Receptor Inhibits Pancreatic Ductal Adenocarcinoma. CMGH. 2020;10(4).

11. Zhu G, Huang Y, Wu C, Wei D, Shi Y. Activation of G-Protein-Coupled Estrogen Receptor Inhibits the Migration of Human Nonsmall Cell Lung Cancer Cells via IKK-beta/NF-kappaB Signals. DNA Cell Biol [Internet]. 2016;35(8):434–42. Available from: https://www.ncbi.nlm.nih.gov/pubmed/27082459

12. Fabian M, Rencz F, Krenacs T, Brodszky V, Harsing J, Nemeth K, et al. Expression of G protein-coupled oestrogen receptor in melanoma and in pregnancy-associated melanoma. J Eur Acad Dermatol Venereol [Internet]. 2017;31(9):1453–61. Available from: https://www.ncbi.nlm.nih.gov/pubmed/28467693

13. Liu Q, Chen Z, Jiang G, Zhou Y, Yang X, Huang H, et al. Epigenetic down regulation of G protein-coupled estrogen receptor (GPER) functions as a tumor suppressor in colorectal cancer. Mol Cancer [Internet]. 2017;16(1):87. Available from: https://www.ncbi.nlm.nih.gov/pubmed/28476123

14. Muller C, Brown-Glaberman UA, Chaney MF, Garyantes T, LoRusso P, McQuade JL, et al. Phase 1 trial of a novel, first-in-class G protein-coupled estrogen receptor (GPER) agonist, LNS8801, in patients with advanced or recurrent treatment-refractory solid malignancies. Journal of Clinical Oncology. 2021;39(15_suppl).

15. Muller C, Chaney MF, Cohen JV, Garyantes T, Lin JJ, LoRusso P, et al. Phase 1b study of the novel first-in-class G protein-coupled estrogen receptor (GPER) agonist, LNS8801, in combination with pembrolizumab in patients with immune checkpoint inhibitor (ICI)-relapsed and refractory solid malignancies and dose escalation update. Journal of Clinical Oncology. 2022 Jun 1;40(16_suppl):2574–2574.

16. Hsu LH, Chu NM, Lin YF, Kao SH. G-Protein Coupled Estrogen Receptor in Breast Cancer. Int J Mol Sci [Internet]. 2019 Jan 2 [cited 2023 Jan 30];20(2). Available from: https://pubmed.ncbi.nlm.nih.gov/30646517/

17. Lappano R, Pisano A, Maggiolini M. GPER Function in Breast Cancer: An Overview. Front Endocrinol (Lausanne) [Internet]. 2014;5:66. Available from: https://www.ncbi.nlm.nih.gov/pubmed/24834064

18. Wei W, Chen ZJ, Zhang KS, Yang XL, Wu YM, Chen XH, et al. The activation of G protein-coupled receptor 30 (GPR30) inhibits proliferation of estrogen receptor-negative breast cancer cells in vitro and in vivo. Cell Death Dis [Internet]. 2014;5:e1428. Available from: https://www.ncbi.nlm.nih.gov/pubmed/25275589

19. Weissenborn C, Ignatov T, Poehlmann A, Wege AK, Costa SD, Zenclussen AC, et al. GPER functions as a tumor suppressor in MCF-7 and SK-BR-3 breast cancer cells. J Cancer Res Clin Oncol [Internet]. 2014;140(4):663–71. Available from: https://www.ncbi.nlm.nih.gov/pubmed/24515910

20. Weissenborn C, Ignatov T, Ochel HJ, Costa SD, Zenclussen AC, Ignatova Z, et al. GPER functions as a tumor suppressor in triple-negative breast cancer cells. J Cancer Res Clin Oncol [Internet]. 2014;140(5):713–23. Available from: https://www.ncbi.nlm.nih.gov/pubmed/24553912

21. Sjöström M, Hartman L, Grabau D, Fornander T, Malmström P, Nordenskjöld B, et al. Lack of G protein-coupled estrogen receptor (GPER) in the plasma membrane is associated with excellent long-term prognosis in breast cancer. Breast Cancer Res Treat [Internet]. 2014 [cited 2023 Jan 30];145(1):61–71. Available from: https://pubmed.ncbi.nlm.nih.gov/24715381/

22. Martin SG, Lebot MN, Sukkarn B, Ball G, Green AR, Rakha EA, et al. Low expression of G protein-coupled oestrogen receptor 1 (GPER) is associated with adverse survival of breast cancer patients. Oncotarget [Internet]. 2018;9(40):25946–56. Available from: https://www.ncbi.nlm.nih.gov/pubmed/29899833

